# *Chloe*: Flexible, Efficient Data Provenance and Management

**DOI:** 10.1101/2020.01.28.923763

**Authors:** Toni Kazic

## Abstract

Reproducible and sharable research requires robust data provenance during and after the experimental process. Each laboratory and experiment has its own goals and methods, and these change frequently. Planning, managing, and collecting data from research crops are particularly labor-intensive tasks, given the tightly compressed time schedule and the operating environments. Moving from a lab’s present record-keeping approach to an electronic ecosystem that improves provenance is an additional burden for groups without dedicated, consistent computational support to make that transition and then to adapt the system as needed. This high barrier to entry and the press of field work makes it easy to postpone “computerizing”.

I have developed *Chloe* to reduce manual effort during experiments and maintain data provenance. A flexible, modular system, *Chloe* integrates simple equipment, data collection strategies, and software into workflows. The design lets one use parts without deploying the whole. This reduces the barriers to entry while still improving workflow efficiency and making *Chloe* accessible to a wide range of users. I offer guidance on ways to adapt *Chloe* to one’s own experimental situation. *Chloe* has been tested and refined with many changes of students, hardware, and experimental goals over the last fourteen years. Though originally designed for maize genetics and computational experiments, *Chloe* can accommodate other types of experiments, wetbench work, and other crops.

## 2 Introduction

Every laboratory produces scientific artefacts with workflows. Whether these are the equations and proofs of a theoretician, the models and databases of a computational scientist, or the data and physical materials of an experimentalist, all travel from the mind of the investigator to archival publication, deposit, and storage. Tracking the provenance of these artefacts over the investigator’s scientific career is an essential first step in making research reproducible and shareable, the goal of the widely endorsed FAIR guidelines [8,19]. Field experiments are particularly challenging for data provenance as the volume of work is high during a compressed time interval and the physical environment can be exhausting. With less reserve capacity, efficiency and reliability in optimizing workflows are especially more important for smaller groups. Small laboratories can now easily generate big data — large volumes of complex, multidimensional data — during a single experiment, but usually don’t have the same access as large groups, consortia, and industry to off-the-shelf laboratory information management systems or dedicated programming support for a custom solution. Finally, any system for provenance must allow investigators to easily adapt it as their needs change while still preserving efficient use of older data and ideas.

Some years ago I suggested ten simple rules for assuring the provenance of experimental data [8]. The rules and their illustrations were based on my experience in an ongoing study of maize disease lesion mimic mutants’ phenotypes, with field, laboratory, and computational experiments. This paper presents the reification of those rules as *Chloe*: a set of design principles; strategies and simple equipment for data collection and management using commodity mobile hardware and services; and open source code for experiment planning, execution, and management. *Chloe* focuses human attention where it matters most — ensuring data quality — by minimizing manual data recording and balancing manual inspection and correction of data with automated data management. All data are collected electronically on mobile devices, simplifying management and processing. The software, protocols, and equipment form an integrated, robust set of workflows built from separable, easily modified components. The system has evolved through multiple changes in hardware, experimental design, and procedures, helping to enforce a reasonable flexibility. Though designed to serve my own experimental needs, the techniques described here should be generally useful and readily adapted for your experiments.

Several principles were of paramount importance when designing and implementing *Chloe*. Chloe is one of the many titles of Demeter, the Greek goddess of agriculture, and means “young green shoot”: I developed *Demeter* before *Chloe*, and use *Chloe* for both the whole system and its Perl component.

- **The computational procedures** should assure provenance; make experimental operations more efficient and robust to error; seamlessly integrate with the physical equipment and procedures used to collect data; provide ample redundant information for error detection and correction; and facilitate information sharing.
- **The data should self-identify and be instantly legible to the naked eye as plain text**, with unique, easily parsed identifiers for each type of artefact. These form the primary keys of the *Demeter* database. Incorporate identifying information into labels, images, audio recordings, files, and filenames. Don’t clutter labels.
- **Make it easy to spot and correct errors**. We prefer Code128 1D to any of the 2D barcodes because it’s easier to see where the barcode is damaged and aim the scanner around that; because they include the identifier in plain text for manual entry when scanning fails; and because the rest of a plant’s information should not be embedded in the identifier since information can change with time and circumstance. Scanning a datum into the wrong column of a spreadsheet is easily detected by looking at the identifier’s prefix, even for pollinations^1^. We strongly prefer interfaces that let us look at multiple screens at once, rather than switching among windows.
- **Only essential data correction should occur in the field or at the bench**, ideally contemporaneously with their gathering. The rest should occur in comfort. Check ambiguities promptly, support multiple collections of the same data instead of forcing their suppression or combination, and minimize manual operations.
- **The code** should rely on free, open-source language implementations and packages; easily port to commonly used platforms and interface to other computational technologies; be fast and flexible enough to accommodate a career’s worth of changes; be reasonably transparent to non-specialists; and facilitate the computation of metadata. Throughout, data and code are processed and edited as plain ASCII text files rather than embedded in more elaborate computational machinery such as relational databases, integrated development environments, or customized graphical user interfaces.
- **Use familiar and well-designed interfaces, hardware, and services whenever possible**. Commodity hardware, software, cloud services, and stationery should have large market shares and long lifetimes. User interfaces should already be familiar, rather than learning a custom GUI. Custom equipment should be simple, directly support data collection and their computations, be easily reused and biodegradable, and be cheap to build and maintain.
- **All components should be mutually independent** as far as possible: *e.g.*, data collection, transfer, cleaning, processing, and database entry are separate from each other and from the generation of labels, tags, pedigrees, and the field book. Use what you need and toss the rest.
- **KISS**: keep it simple, stupid.

## 3 Materials and Methods

### 3.1 Making *Chloe* Work for You

Before the trees, the forest: how hard is it to adapt *Chloe* to the needs of one’s own group?

- **Equipment** Beyond your favorite mobile device, the only commodity equipment you don’t already have is probably just a stand-alone, matchbox-sized bluetooth 1D barcode scanner. We build and maintain our barcoded row stakes in the laboratory (Section 3.3). If you decide to tag plants, your machine shop can easily build the fixture used in their manufacture and the pins and blocks used to organize them (Section 3.2).
- **Procedures** The three main changes are scanning into spreadsheets or taking notes electronically instead of writing on paper; setting up per-plant or per-row data collection by tagging; and maintaining barcoded labels. Electronic notes include audio recordings of in-field commentary. Tagging is postponed for as long as practicable to minimize shifting awkwardly placed tags. Tags for stocks with unreadable parental barcodes are reprinted and stapled over the originals as these are discovered during shelling and packing.
- **Code** Three parts of *Chloe* and *Demeter* must be adapted for your organization: your laboratory’s names, locations, directory trees, and seed filing order; construction and parsing of identifiers; and which data you collect. These are grouped in separate modules for easier modification. The rest of the computational infrastructure is either free and platform-independent, or free but platform-dependent, or cheap and platform-dependent.

### 3.2 Materials

#### Computational Languages

*Chloe* is written in Perl, version 5.26; *Demeter* is written in SWI-Prolog, version 8.0.3. Both are free, run on any platform, have excellent and extensive tutorials, and are fully documented. *Chloe* and *Demeter* should be compatible with older versions of these languages, and forward compatibility is highly likely as the use of external modules has been minimized. Throughout, I use the following notational conventions. Identifier elements, commands, Prolog rules, and filenames are in `this monospace font`. Prolog files are named with the .`pl` extension and Perl files with the .`perl` extension. This is consistent with Prolog, but not Perl, practice. By convention, names of Prolog rules include the number of arguments (*e.g.*, `foo/3`).

#### Computational Packages

The rest of the computational infrastructure is either free or very inexpensive, and versions of all of these packages exist for all platforms. *Chloe* assumes you have GNU Barcode, LATEX, ImageMagick, and GNU enscript installed. For editing files, I strongly recommend Emacs and its included org mode. Other suitable plain ASCII editors include TextEdit on the Mac and NotePad on Windows (be careful to save as plain ASCII!), and vim on Linux and Mac — but not for example Word, Pages, or LibreOffice Writer, which embed troublesome formatting information in the output file. We use the beorg note-taking app in the field, which offers a subset of org mode and is cloud-compatible. We use Apple’s Numbers spreadsheet to collect data. There are many choices in spreadsheet applications: not all will share nicely with one’s preferred cloud service, easily timestamp an entry, or leave earlier timestamps unchanged as new ones are entered. Similarly, there are multiple choices for cloud services and pdf annotation apps. We use Dropbox, which interacts well with our other apps. We prefer iAnnotate to Adobe Reader because pdf annotation and cloud syncing are more robust and versatile. Installation and configuration instructions and workflow details are in *Chloe*’s documentation on GitHub.

#### Equipment

As Figure 1 shows, most of *Chloe*’s equipment is pieces of paper. We code plants for lesion phenotype, pollinations, and photography using cut 9 inch colored paper twist ties (Bedford Industries). We use Apple’s iPhones and iPads running the data collection workbook to collect data using a KoamTac KDC200i matchbox-sized bluetooth 1D barcode scanner. These are available in versions compatible with other operating systems. We find it easier to use a scanner physically separated from the mobile device, rather than forcing the same person to read and record the data or to toggle between scanner and spreadsheet apps.

**Figure 1:**
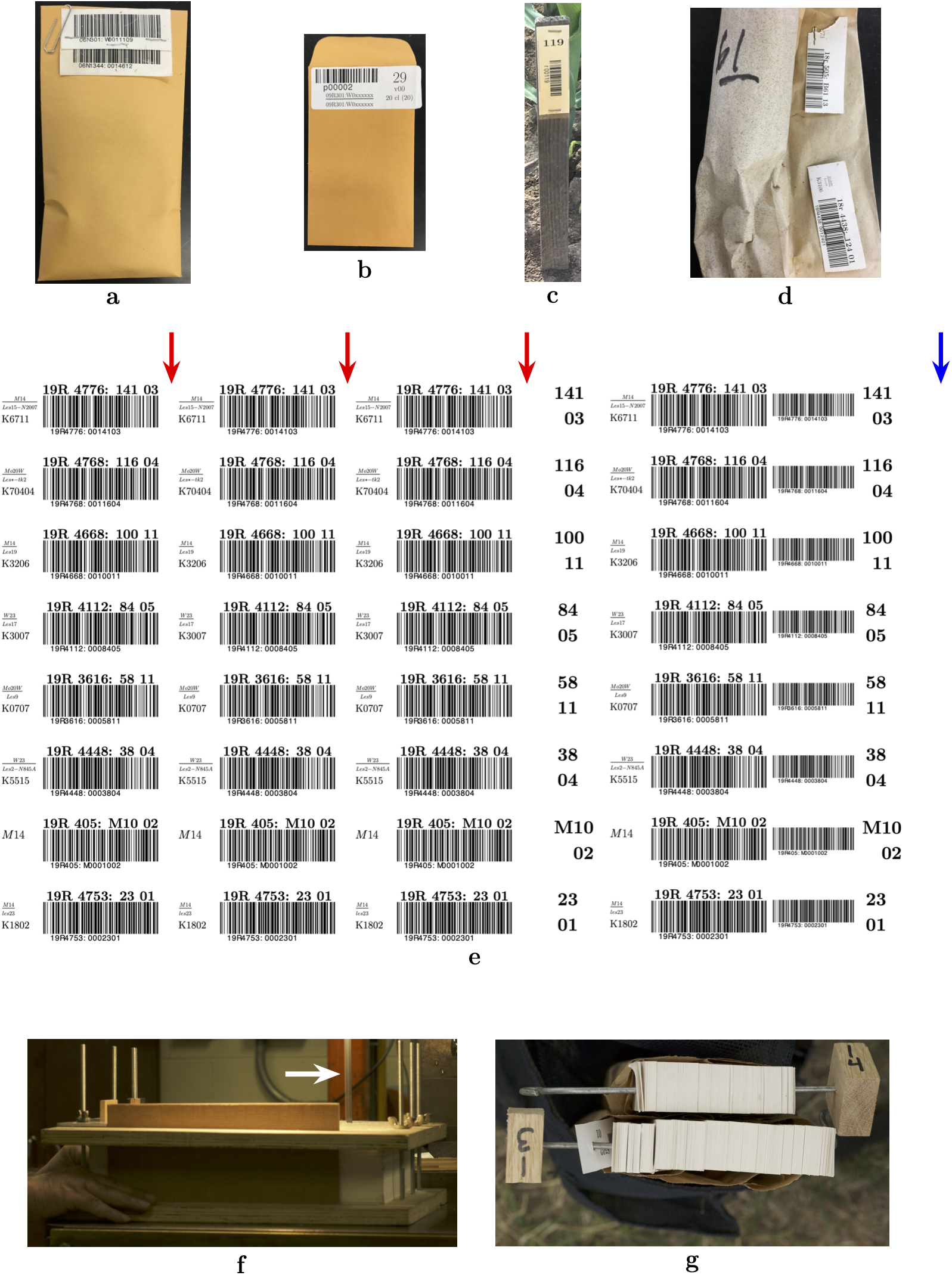
*Chloe*’s equipment (not to scale). (a) A seed storage envelope, with the maternal tag stapled above and to the paternal tag. (b) A seed packet labelled with a barcoded label. (c) A row stake in action; this one is at least two years old. (d) A pollination bag with the maternal tag above the paternal. These tags are removed and stapled to the seed storage envelope at shelling. (e) A sheet of plant tags. The bottom tag begins the first bundle and consecutive tags are laid out down the stack of tags first, then across bundles. Approximate locations of perforations (red arrows) and the pin holes (blue arrow) are shown. (f) Sawing apart the tags on the band saw. The drilled block of tags is clamped in the plywood fixture and stacks are sawn individually. White arrow marks the saw blade. (g) Tag bundles in the left apron pocket with galvanized pins and wooden blocks. Bundle 13 is ready for tagging; tags will come off to the left after the block is removed.

Seed stocks are stored in no. 7 coin envelopes (3.5 × 6.5 inches); for planting, seed is packed in no. 3 coin envelopes (2.5 × 4.25 inches) labelled with a 1 × 2 5/8 inch Avery label. Plant tags are printed on tabloid-sized 100 lb white card stock (11 × 17 inches, not weather proof). The unfinished wooden row stakes are 18 × 11/8 × 1/4 inches (Hummert); their labels are printed on ordinary printer paper and laminated.

For drilling and sawing blocks of tags, our machine shop built a plywood fixture. This is two sheets, approximately 18 × 11 × 3/4 inches, drilled to admit three pairs of 12 inch carriage bolts to clamp the block of tags in place. Vertical stops are set in slots along a long and a short edge, and the other long edge is cut to admit the saw blade. Pins for bundles of tags are made from no. 9 galvanized wire, bent at one end and sharpened at the other; the wooden blocks are sawn from 1 × 1 × 1/2 inch oak.

A simple “counting pan” was routed from 14 × 18 × 1/2 inch pine. This is ten round-bottomed spouted grooves, spaced about 2 inches apart, with lines drawn at intervals of five large kernels. The grooves are painted with black Sharpie for better contrast and the pan is set on a piece of rubbery shelf liner to prevent wiggling during kernel sprinkling and sweeping. Ears are shelled using a new device that rotates the ear against shelling flutes. Kernels drop into a mesh tray (positioned over a trash can) that is shaken once for final cleaning, then poured into the prepared seed stock envelope (McInturf *et al.*, in preparation).

### 3.3 Methods

#### Data Collection

Data are collected into spreadsheets carried on the mobile devices using the bluetooth scanner, selecting menu options, and timestamping with a datetime function. Direct typing is minimized as much as possible. The spreadsheets’ columns are ordered left to right in decreasing frequency of being changed and prefilled with relatively unchanging data as much as possible. The last column is always named by the type of datum in the spreadsheet, but is otherwise unused. A master workbook with all the data collection spreadsheets is revised before each crop to reflect changes in experimental goals, methods, and personnel. The name of each spreadsheet begins with `raw`. to so identify its contents in its exported `csv` file. Revisions are checked against *Chloe*’s data processing scripts to ensure the latter anticipate the current column order and data formats. Changes during the season, for example to capture new ideas, experiments, or types of data, are incorporated in the same way in the workbook, *Chloe*’s data conversion scripts, and *Demeter*’s database design. The master workbook and convenient subsets of it are maintained in a directory in our cloud service and also in a corresponding directory in our main file tree.

On data collection days, a subdirectory with the date is created in the cloud and the appropriate workbook copied to it, changing its name to include that day’s day and month and deleting any spreadsheets that won’t be used. Consistent with *Chloe*’s data transition model, the filename change signals its contents will be the raw data (see Figure 3). Each mobile device has its own branch of the crop’s data directory tree so that multiple devices can be used on the same day with the same file naming scheme. Once a copy of the day’s workbook has been downloaded to a device, placing the device in airplane mode avoids time-consuming synchronizations in the field. During breaks and at the end of the day, we reconnect the device to the cloud and export the day’s workbook as a bundle of `csv` files, letting the cloud service synchronize them to the day’s cloud subdirectory on our laboratory computers. By default, a datum need not be unique in *Chloe* and *Demeter*, thus supporting temporal changes, differences of opinion among observers, and simply forgetting what was last done.

Each datum is timestamped as it is collected, but the length of the interval between action and timestamping varies by the type of datum. The importance of temporal precision and the ease of data collection determine the length of this interval. For example, packing a line’s seed, planting a row, setting up tassel bags, pollinating, tissue collection, and photography are timestamped within a minute or two; cutting back ears are timestamped within four–five hours; and mutant scoring and harvesting are timestamped within a day. We score phenotypes of interest as each row is ready, marking lesion mimic mutants with twist ties looped around the culm at a convenient place and twisted together twice to lock the loop. The scoring date of each row is written on a shoot bag stapled around the first tall plant in the row. These data are collected later, usually after pollinations are complete. Some operations are not timestamped at all, reflecting consistency in our procedures or choice of interesting details. For example, *ex situ* leaf photography involves cutting, rinsing, and air drying leaves before photography, and we stage the work so that no more than 90 minutes elapses between collecting the first leaf and imaging the last. Similarly, we do not record dates of field management tasks, such as irrigation and hoeing. These choices will vary by laboratory, but should be fairly consistent within a lab.

#### Seed Management

Packet labels are generated with a packet identifier, parental numerical genotypes, seed location, and packing and planting instructions. These are sorted into seed inventory filing order by `pack_corn/1`, which then calls `make_seed_packet_labels.perl` to print them on Avery labels to form a packing checklist. Inbred, elite, and “packets” for rows left unplanted for image processing reasons have standard packet identifiers, while unique packet identifiers for each packet of mutant lines are generated. As seed is packed, the label is applied to the packet, and both it and the parental numerical genotype labels from the seed stock are scanned and timestamped. Packets are filed for planting in row sequence order as they are packed. The counting pan is used when many packets of the same line are needed. Seed is sprinkled from the stock into a groove, the excess is pushed away from the spout using the lines to visually count kernels, and the seed swept into the labelled packet. Excess is swept back into the seed stock. The pan can be used for single packets of a line, but we pack only one line at a time to prevent contamination.

`make_harvest_plan.perl` computes 40 days after the last pollination in each row and groups rows for harvest by planned harvest date. Ears are harvested into their pollination bags after brief cleaning in the field. Each row stake is scanned as the row is harvested to timestamp the harvest (`row_harvested/5`). Our shelling workflow integrates data collection, shelling, seed filing, and inventory management. Prior to shelling an ear, the parental tags on the pollination bag (Figure 1, panel d) are scanned, the number of kernels estimated or counted, any notes or comments on the ear entered, and the tags removed and stapled to each other and then to the seed stock envelope (Figure 1, panel a). If either parental tag is too degraded to scan, the tag is entered in the spreadsheet as a rowplant to record this situation and saved separately; the data are then cleaned and pushed to *Demeter*. Stock tags to replace degraded parental tags are generated with both parental barcodes by `make_new_seed_labels.perl`, printed on card stock, cut, and stapled over the originals when packing the next crop or when packing that line for planting, whichever comes first. After all the ears are shelled, the maternal tags of the first and last ear in each sleeve and the sleeve barcode are scanned and timestamped. These and the `row_harvested/5` data are incorporated into the `harvest/6` and `inventory/7` database facts using `update_inventory.perl`. A running inventory is kept so that refiling seed stocks or a stock’s significant depletion can be recorded as needed. The crop planning code uses only the most recent inventory datum for a stock. The filing order for our seed inventory is: crop year; nursery location; inbred lines; mutant lines. Within each category of lines, seed is filed in rowplant order in virtual “sleeves” formed by wooden dividers in the cardboard seed boxes; the sleeves are barcoded. `scootch_sleeve_bdries.perl` adjusts `inventory/7` facts if the corn is refiled.

#### Row Stakes

Rows are numbered across all fields when multiple fields are planted, so that each crop has only unique row identifiers. Row labels are printed on ordinary paper, laminated, and then sliced apart with a paper cutter without bothering to seal the sliced edges. The label is stapled near the top of the stake with galvanized staples, pounding in the protruding tips to prevent injury. The stakes are rubber-banded together in bundles of 20 and repacked in the boxes the stock came in, ready for planting. With laminated labels, the stakes last at least several years before they become inconveniently short; rot is minimized by inserting the stake in the soil to about 1.5 inch depth. After harvest, stakes are collected, excess soil scraped off against plants, packed loosely in numbered mesh bags, hosed off, and dried before packing them away for next season. When a stake has passed its useful lifetime, the label is torn off for separate discard and the stake tossed into the field to decompose.

#### Planting and Stand Counts

Seed is planted by hand for mutant and inbred lines, where stand count information is important; or using a one-row Jang TD1 mechanical push planter (Woodward Crossing) for the border and rows planted with elite lines, where stand counts are not taken. The barcodes on the packet and row stake are scanned as each row is about to be planted for all but the border rows. For stand counts, the row stake is scanned into the spreadsheet and the rest of the data entered before timestamping the datum.

#### Plant Tag Manufacture, Placement, and Use

A sheet of tags is shown in Figure 1, panel e. The crop’s planning, genotype, planting, and stand count data are compiled with either Prolog (`generate_plant_tags_file/3`) or Perl scripts to tabulate the information for plant tag production (the Perl code needs some human supervision). This rule calls `make_plant_tags.perl`, which generates the plant tags’ pdf file. Laying out eight tags/sheet, the algorithm stacks the tags so that the next tag in a row is in the same column on the sheet below its predecessor, not on the same sheet. Our plant tags include three barcoded tear-off tags for marking pollination bags and leaves for photography, two additional barcodes, basic genotypes, and rowplant numbers in large font.

A commercial firm (FedEx) prints the tags on tabloid-sized card stock (11 × 17 inches), cuts away the excess paper to leave a legal-sized 8.5 × 14 inch sheet, and perforates the tear-offs with a tooth spacing of approximately 1 mm (Figure 1, red arrows in panel e). Cutting away the excess eliminates fussy alignment of the page of tags to smaller sheets. Our machine shop then places the block of sheets in the clamping fixture (Section 3.2), drills a hole near the edge for each stack of tags (blue arrow, Figure 1, panel e), and saws them apart on a band saw (Figure 1, panel f). As each stack is removed from the fixture, a pin is inserted in the hole and then stopped with a numbered wooden block. Each bundle of tags is rubber-banded, placed in a numbered tassel bag, and rubber-banded again. The pin and stop prevent accidentally disordering the tags and let one easily transfer tags among bundles so that each bundle begins and ends at a row, if desired. To tag, the bundle is fanned, and returned to the now loosely rubber-banded tassel bag, the block removed, and the bundle placed in a deep apron pocket (panel g). During tagging, the tags are slipped off the pin, looped around the plant, and stapled near the top to span the first tear-off perforation so that water absorption does not cause the tear-offs to separate from the loop.

Our pollinations are individual by plant, our lines differ considerably in their maturation rates, and a single male can be used several times on different maternal parents: so plants used as males and females must be individually tracked. Plants likely to be used as males are marked with a twist tie and a tear-off tag collected. These tags are organized to simplify monitoring the tassels for flowering. When the tassel is bagged to collect pollen, the bag is labelled with the tear-off tag, scanned, and timestamped. A tear-off is collected when an ear’s silks are cut for later scanning. These data are recorded in `cross_prep/5` and used to plan the next day’s pollinations. When an ear is pollinated, its tear-off tag is stapled to the bag above the male tag (Figure 1, panel d).

#### Pedigree Construction

*Demeter*’s pedigree computation exploits the provenanced data collected over all the crop cycles. The `build_pedigrees/2` rule recursively computes all descendants from the founding lines using the numerical genotypes of both parents of successful pollinations. The founding lines are defined by the `founder/9` rule described in Section 4.2. At each step in the depth-first search through the descendants, determining if a pollination recorded in the cross/8 facts was successful uses the `harvest/6` facts, which include the parents’ numerical genotypes and an approximate or actual kernel count made during shelling. Building the numerical genotypes of the parents relies on the data in `packed_packet/7`, `planted/8`, and the most recent stand count in the `row_status/7` facts. The family numbers in the numerical genotype are automatically assigned when a line is first planted, producing a new `genotype/11` fact that includes a proposed symbolic genotype that is manually checked for correctness. `packed_packet/7` data includes scans of the (now grand)parental numerical genotypes on the seed storage envelope during packing, and `planted/8` includes scans of the barcoded seed packets and row stakes. The output includes the path to the image of a mutant parent’s leaf (data from the `image/9` facts). For convenience, we have organized our lines into trees by the initial mutation of interest (stored in `pedigree_tree/2`). The names of the output files are generated from the lower-cased mutant names with Unix special characters, such as ‘*’, ablated. Both plain text and pdf versions of each pedigree are placed in their corresponding subdirectories defined by `pedigree_tree/2`. A copy of the pdf file tree is placed in our cloud service to facilitate annotation on a tablet. After backing them up to our laboratory and cluster storage, the annotated files remain in the cloud throughout the field season for viewing in the field.

### 3.4 Availability

Code, sample data, and documentation are in *Chloe*’s GitHub repository. The documentation includes installation, configuration, and usage suggestions and describes several workflows in detail. *Chloe* is licensed under the terms of the GNU Affero General Public License, which allows one to freely download and modify the code, or to run it on a server, provided one specifies these modifications and uses the same license. Please use GitHub’s issue tracking machinery to submit comments, suggestions, bug reports, and feature requests. Please cite this work as specified in *Chloe*’s README file.

## 4 Tracking and Storing Data Provenance

*Chloe*’s simple equipment is designed for capturing and maintaining provenance. The barcoded labels are produced; applied to the different artefacts; and scanned into the workbooks to collect data. Data move through a set of states to land in *Demeter*. Manual tasks in this process are to position the labels on envelopes, row stakes, and plants; to enter the data; and to check, clean if necessary, and copy the data before processing.

### 4.1 Data Movement in *Chloe*

#### Classes of Experimental Data and Data Flow

The field component of our research on complex phenotypes mixes the creation of near-isogenic lines of maize disease lesion mimic mutants, phenotypic observations and photography, and tissue collection. The field worklow has four classes of data, all collected and used on mobile devices:

- tabular data collected by scanning barcodes and punching data into spreadsheets, processed and deposited in *Demeter*;
- pdf files of the pedigrees and the field book. The field book is annotated throughout the growing season: new notes are incorporated into *Demeter* and the field book recomputed as needed;
- images and audio recordings collected on either HD storage cards or on a mobile device and directly transferred to laboratory and cluster storage; and
- ephemeral notes on tasks that need followup in the field or laboratory, taken and maintained in a cloud-aware note-taking application.

Figure 2 is an overview of the flow of data among the field and seed room, laboratory computers, *Demeter* and *Chloe* modules, cloud services, physical devices, and the cluster’s storage. Data collection is supported by generating barcoded labels and tags for each type of experimental artefact. *Chloe* has a set of Perl scripts to do this, one for each type of artefact, that call a set of shared subroutines. We now outsource the steps in plant tag manufacture between file generation and plant tagging (see Section 3.3 for details). In addition to standard genetic tasks, other manual operations set up the physical equipment for provenance, such as maintaining barcoded row stakes and tagging each plant used for genetics or phenotyping. Computational tasks are a mix of manual and automated operations. For example, transfer of data collected in the field to the cloud is automated *via* cloud services; error correction is manual; and database entry and pedigree construction are automated.

**Figure 2:**
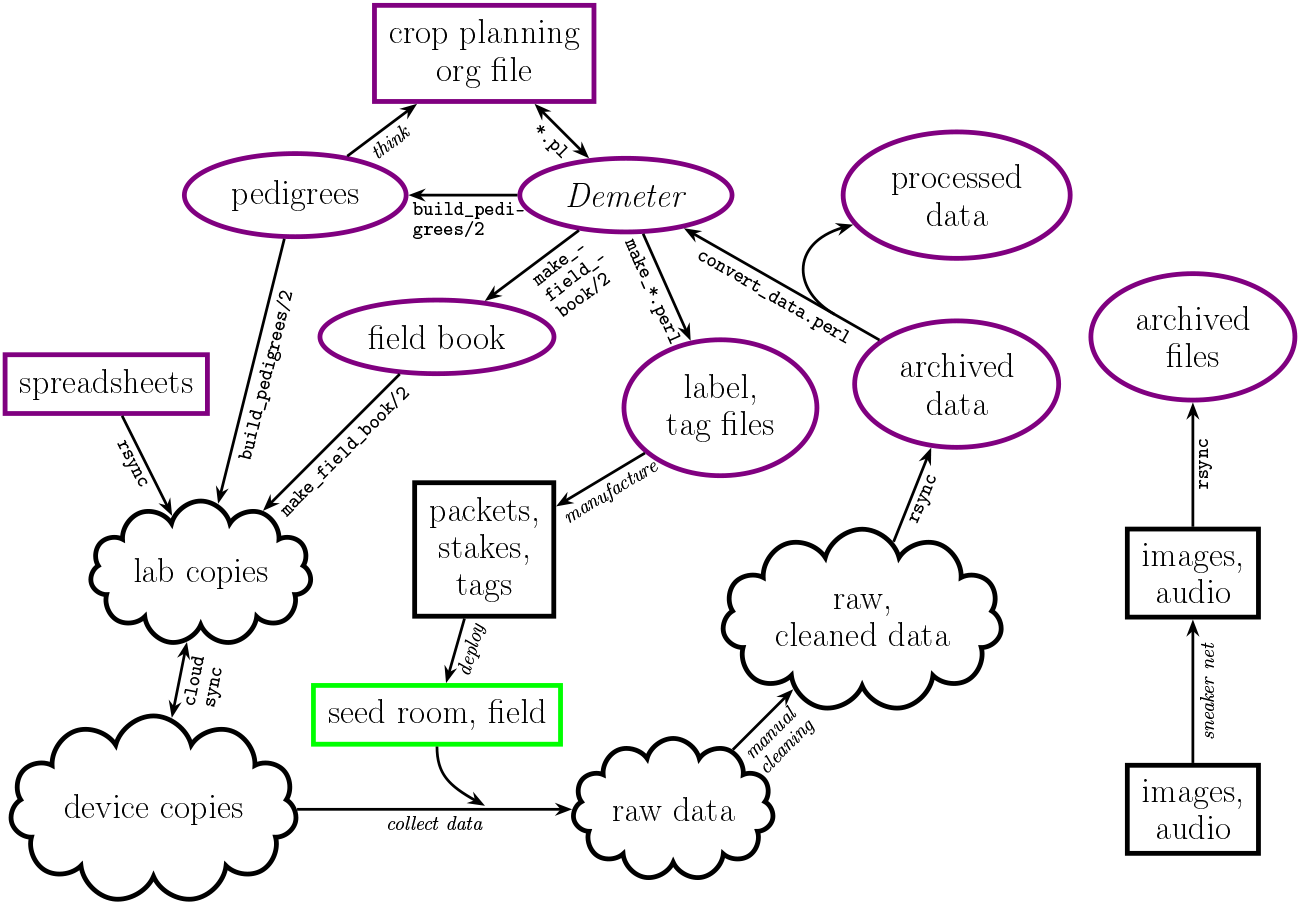
Information flow in *Chloe*. Rectangles denote tasks performed manually; ellipses, the outputs from *Chloe* and *Demeter*; and clouds, well, … cloud services, of course. Names of `scripts, rules, or commands` and *manual operations* describe the main operations in each transformation. All shapes outlined in violet are `rsynced` to cluster storage at least nightly.

#### Identifiers

The heart of provenance tracking is the assignment and use of unique identifiers, barcoded on labels and plant tags to speed collection and self-identification of data. Several identifiers and their encoded information are illustrated in Table 1. The six character identifiers for physical artefacts begin with a unique letter indicating what type of artefact it denotes (a semi-explicit semantics^2^): `x`, seed boxes; `a`, seed bags; `v`, seed filing sleeves; `p`, seed packets; `r`, rows; `e`, tissue samples; and t, greenhouse pots and flats. Denoted by 15 character strings, each plant’s numerical genotype is the primary key in *Demeter*, the experiment database. The numerical genotype identifies each plant in both time and space. All data, field operations, samples, and seed from a plant are referenced in *Demeter* using the numerical genotype, tracking their provenance.

**Table 1:** The two types of identifiers: 15 characters for the numerical genotypes of individual plants; 6 characters for packets, rows, pots, flats, boxes, sleeves, seed bags, and tissue samples. The identifiers’ semantics are described in the text.

|  |  |  |  |
| --- | --- | --- | --- |
| 06R0011:0000000 | family 11 founder: seed from others, or maternal plant in someone else's field | x00000 | box, this one for founders |
| 09R0119:0015403 | plant 3 of founder 119 planted in row 154 in the 2009 summer nursery | v00302 | sleeve in box |
| 07R201:S0xxxxxx | maternal parent of Mo20W pooled inbred seed from sibbing; constituent ears tracked separately | a00001 | bag of pooled inbred seed |
| 07R301:W0yyyyyy | paternal parent of W23 pooled inbred seed from selfing, constituents tracked | p00002 | packet; standard packet number for W23 inbred parent |
| 19R205:S0000103 | Mo20W inbred plant 3 in row 1 of the 2019 summer nursery | e00783 | tissue sample |
| 14R4186:0026908 | plant 8 in mutant family 4186, planted in 2014 summer nursery row 269 | r00269 | row 269 |
| 07G301:W0000806 | W23 parent from 2007 greenhouse nursery; row and plant arbitrary | t00193 | pot, flat, or Petri dish; specified in the `Soil` argument of `planted/8` |
| 06R0000:0000000 | empty seed for skipped row | p00000 | empty packet filled with empty seed |

Referring to the examples in Table 1, the numerical genotype includes the year the crop was planted (`06,07,09,14,19`); a nursery identifier (summer, winter, greenhouse, `R,N,G`, respectively); the family number (`0011,0119,201,4186`, *etc*.) and if a recurrent inbred parent or elite line, a family letter (`S,W; M,B,L for M14, B73`, and an elite line, respectively, are not shown); row (`00000,00154,0xxxx,00001`, *etc*.); and plant (`00,03,yy,06,08`, *etc*.). Though not essential, letting each type of artefact and line have a unique case-insensitive, at least partially mnemomic, letter is very helpful in immediately spotting scanning errors and confusions.

#### State Transitions of the Data

As they are collected, saved, and processed, data transition among a set of states. Progress through these events is tracked with the simple state transition model shown in Figure 3, in which the state of a file’s data is indicated by a prefix to the file’s self-identifying name. This approach decouples data collection in the field from their manual cleaning and automated processing. Our experience is that contemporaneous cleaning and processing is optimal, provided one is not exhausted. Having those who collect the data clean them tends to improve the quality of subsequent collections.

**Figure 3:**
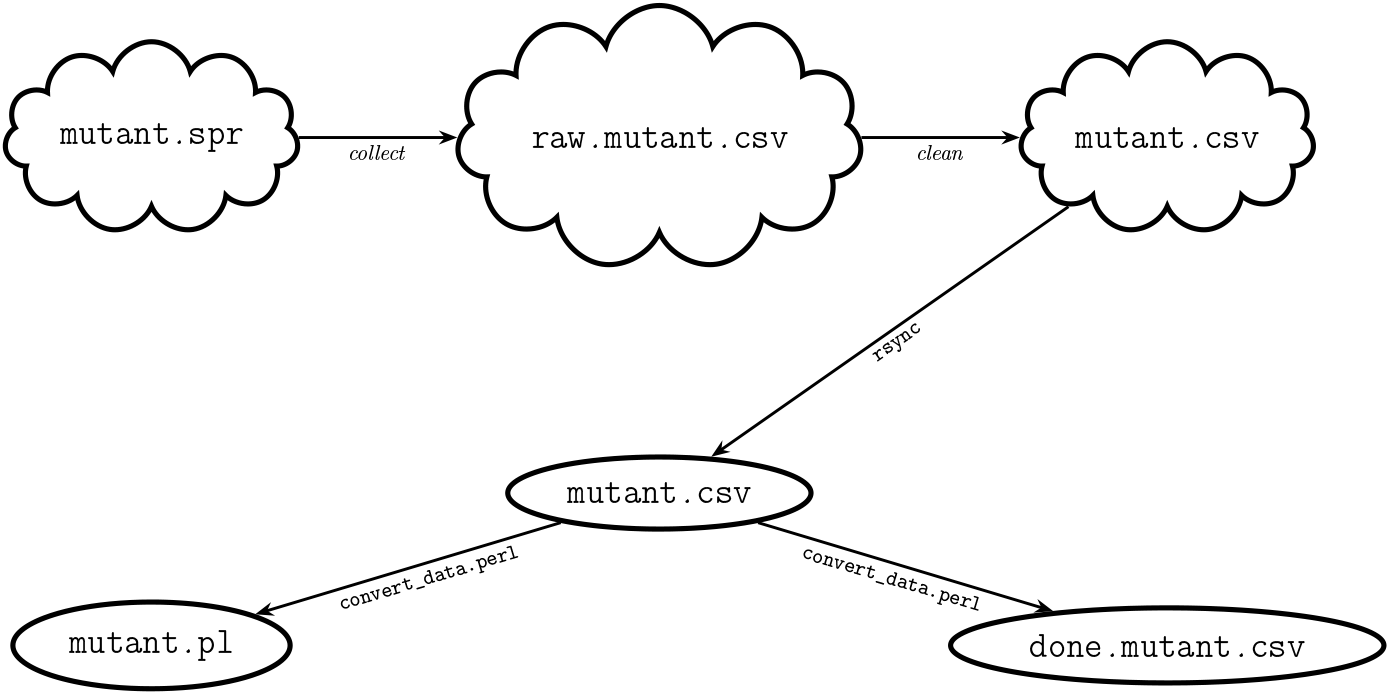
Transitions in the state of *Chloe*’s data, using mutant phenotype information as an example. Transitions are marked by filenames and locations (clouds for cloud services, ellipses for local and cluster storage). Each spreadsheet and `csv` file is in a subdirectory named by the date of data collection (not shown). `spr` denotes a generic spreadsheet in a workbook.

When data collection is finished for the day, we unzip the csv bundle and manually correct any errors in the exported `csv` files in the cloud directory. The `raw`. prefix is dropped from the final versions of the data files, signalling these are cleaned data ready for processing by *Chloe*: the original raw data files are retained as either zips or the completed workbook in its native file format. The cloud subdirectories are then `rsynced` to the unprocessed data subdirectories on our laboratory computers. *Chloe* includes a set of Perl scripts that process the cleaned data and deposit them in the *Demeter* database, renaming successfully processed files with the prefix `done`. to prevent further processing. A master script, `convert_data.perl`, takes as inputs the day and month of data collection and a processing option (letting one check the output data before committing them to the database) and calls members of a set of spreadsheet-specific data processing scripts based on the name of the last column of each `csv` file. Database deposit can be reversed by deleting or commenting out bogus data in the *Demeter* file, renaming `done.*.csv to *.csv`, and repeating processing. Figure 4 shows the key elements of this process using mutant scoring data as an example. We preserve the raw data, their manually cleaned version, and the processed version indefinitely.

**Figure 4:**
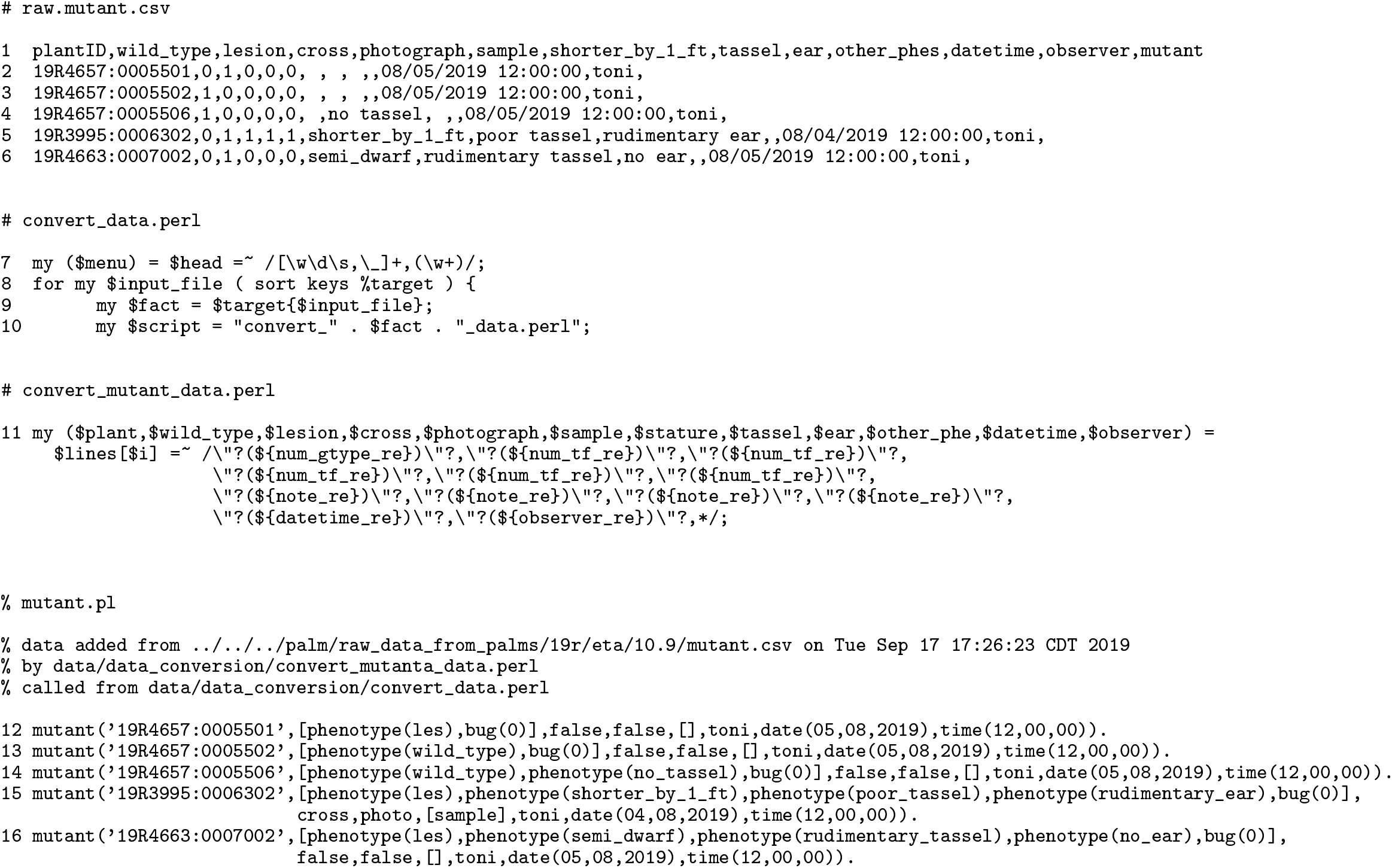
Transforming raw data into *Demeter*; line numbers are in the left-most column. (top) Spreadsheet header and selected data from the `raw.mutant.csv` file (lines 1–6). The scoring date was August 5 (written on a shoot bag stapled to the first plant in each row), but the data were collected on September 10. Column formats include text, numeric, datetime, and drop-down menus for relative plant height and reproductive organs. The last header field identifies the type of data. (middle) Key extracts from `convert_data.perl` and `convert_mutant_data.perl`. The first part grabs the name of the menu from the head of the file and constructs the name of the script that will be called to process the mutant data (lines 7–10). The regular expression (line 11) in `convert_mutant_data.perl` parses the cleaned `mutant.csv` file and stores the data for further processing: the component regular expressions are in `label_making/Typesetting/MaizeRegEx.pm`. (bottom) The output facts and their header in *Demeter*’s `mutant.pl` file (lines 12–16; intra-fact line feeds added for clarity). The 10.9/`mutant.csv` file has been moved to 10.9/`done.mutant.csv`.

### 4.2 *Demeter*, The Experiment Database

*Demeter* is our declarative database of experimental data. It includes the crop preparation and field data, information on leaf photographs and tissue samples, and a number of computations to help manage experiments. *Demeter* is written in Prolog, a well-known logic programming language used in artificial intelligence applications. Prolog has some important advantages for non-specialists. The data (“facts”) and code (“rules”)^3^ are in plain ASCII that is easily edited in an ordinary text editor; pedigree tracing and other naturally recursive computations are easily specified and efficiently executed; and prototyping databases is straightforward [9].

Most importantly, Prolog is a full, expressive programming language that very naturally captures complex ideas as logical rules, rather than being limited to the computational algebra of relational databases. The rules explicitly describe one’s ideas about the experiments and their underlying biology by specifying the logical properties a successful computation should satisfy, rather than getting lost in procedural thickets. New ideas can be added alongside the old by *intension* — specifying a rule that describes the idea — rather than explicitly declaring each item that one believes meets the rule (*extension*^4^). The evolution of my version of *Demeter*’s `founder/9` rule is a good example. I originally allocated family numbers 1–199 for mutant founding lines and 200–599 for inbred lines, starting my own lines at family 1000. As I acquired more founding mutant lines and landraces, I needed more family numbers that did not collide with the additional categories of lines I had also accumulated that have families between 600 and 1000 — elite lines, sweet corn, popcorn, *etc*. The current version of `founder/9` checks a family number that is less than 1000 against rules defined for each type of maize. Should I someday need founders with numbers above 1000, I’ll just add a new rule for them to `founder/9`, rather than renumber all the families, plants, and their data.

*Demeter*’s rules reify genetic ideas and our procedures for field and data management. These fall into four main categories:

**inference** rules that build pedigrees, choose lines for planting, construct genotypes, and generate the field book;
**monitoring** rules that monitor the state of the field work, such as progress in photography and scoring;
**utility** rules that build and parse identifiers, manage directories and seed inventory, generate indices, print output files, *etc*.; and
**cleaning** rules that detect missing or messy data and help one clean them.

For example, a common genetic task is to construct one or more pedigrees, often as a prelude to other analyses (*e.g.*, reference 17). *Demeter*’s pedigree computation illustrates how a rule can reach across many types of facts as well as over multiple crops. Figure 5 shows an excerpt of a pedigree after annotating the pdf. Briefly, from the founding lines defined by the `founder/9` rule, the descendants are recursively computed using the numerical genotypes of both parents of successful pollinations. For each generation, data from pollinations, harvest, stand counts, planting, and seed packing determine the relationship between the numerical genotypes of parents and offstpring. Auxiliary information, such as the location of images of the male parents’ leaves, are gathered for output. Each pedigree is written to an ASCII and a pdf file, named by the mutation of interest. The pdf files are copied to our cloud service for manual annotation; they remain there throughout the season and are backed up to our laboratory and cluster storage. More computational details are in Section 3.3.

**Figure 5:**
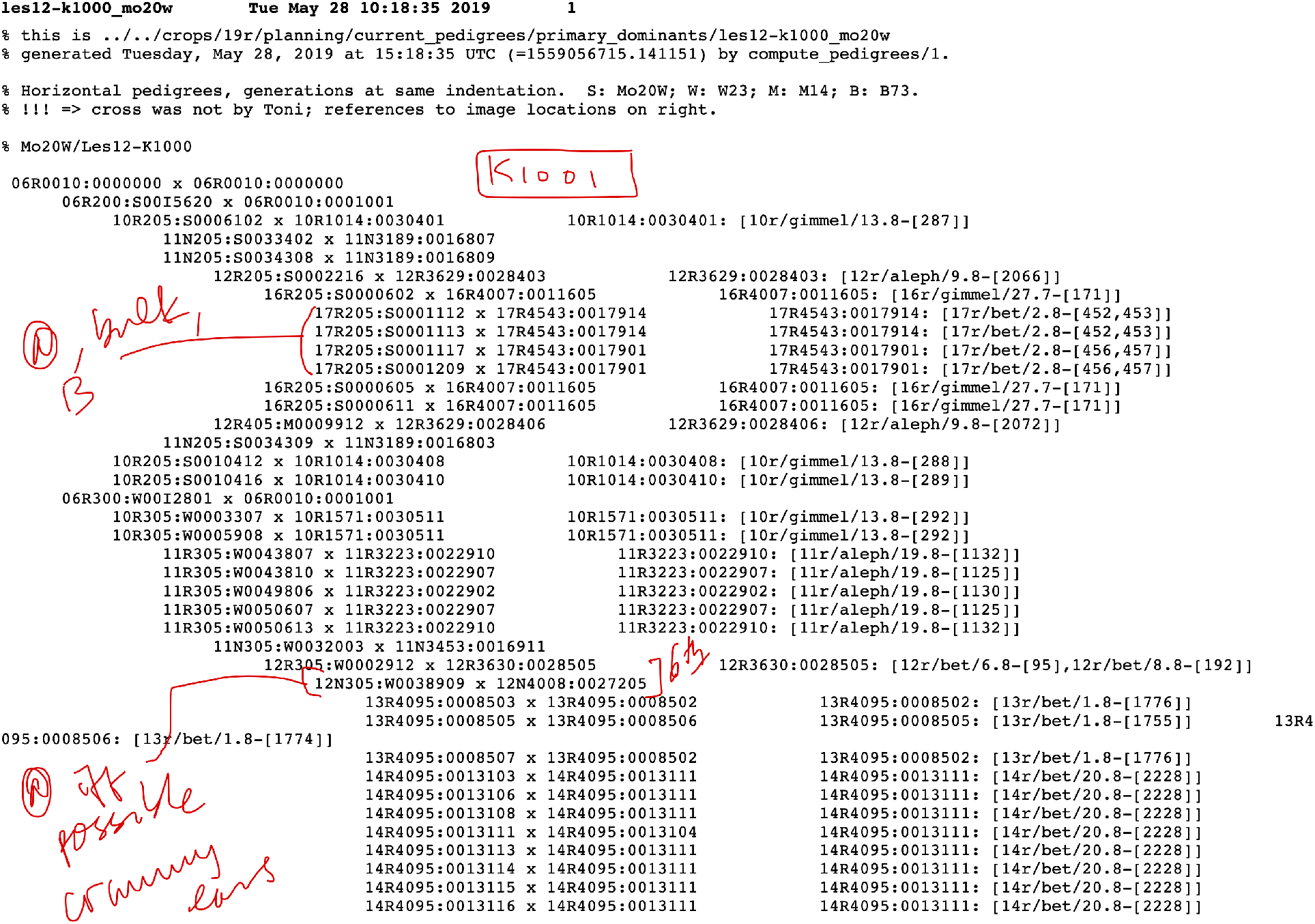
The first page of a typical pedigree file after manual annotation (red). After the file header information, the pedigree is on the left of the page and the path to the parental images are on the right.

On the facts side, a plant’s numerical genotype is the primary key for its information in *Demeter*. Each type of datum has its own file (making the equivalent of a relational database table, but directly editable), and each batch of data begins with a comment describing how and when the data were added to *Demeter*. Data that need to be changed after accession into *Demeter* — for example, because a new mutant has been recognized retrospectively and needs a new family number, or an error in cleaning the raw data was overlooked — are changed in the corresponding *Demeter* file, adding in a comment with the rationale, author, date, and original datum. The file `dem_facts.pl` in *Chloe*’s GitHub repository synopsizes the database facts. Like writing new logical rules at any time, adding new types of data during the field season is as easily done: just adapt a related spreadsheet and its Perl processing script, and create a new Prolog file to receive the data. Some data are deliberately not incorporated directly into *Demeter*, but instead referenced in accordance with common database practice. For example, leaf images are not stored in *Demeter*, but the `image/9` fact stores the numerical genotype, details of the photograph’s composition, image number, camera name, and datetime. The last three, together with the crop identifier extracted from the numerical genotype, are sufficient to immediately construct the relative paths to all images of that plant, as is done in `build_pedigrees/2`. Figure 5 shows these references on the right hand side of a sample pedigree.

## 5 Managing Hiccups and Failures

Nothing replaces good management of the field, and dealing with the inevitable procedural hiccups is part of that. We’ve encountered three main types of problem. The first is an intentional or accidental failure to collect data: a commercial nursery service will not scan packets and stakes when planting, or someone will forget to scan something. Such data can be easily reconstructed by hand from one’s knowledge of the procedures; we insert comments in the files indicating the problem and approach to reconstruction so it is clear those data are not contemporaneous. The second is bad or absent barcodes. The plant tag card stock is not weather-resistant and a few barcodes will degrade. We either type in the rowplant during data collection for later augmentation; or scan the tag for another plant in the row and edit that on the spot; or number a row’s plants with a Sharpie and then photograph hand-written parental rowplants on the pollination bag. The choice of method is governed by whether the barcodes will be needed later. We save the rowplant-only information before correcting it so we have a list for making replacement labels for seed storage. For pollinations, we enter the pollination data from the photographs and their timestamps, then staple at least the maternal tag to the pollination bags, being extremely careful to match the tag’s identifier to the previously assigned rowplant and double-checking the correctness of all data using that identifier.

The syntactic problems just described are easy to detect: *Chloe* and *Demeter* protest when data are missing or incorrectly formatted. In contrast, detecting the third type of problem — wrong semantics — depends heavily on one’s knowledge of the experiment and coworkers and care in managing everyone’s fatigue. Plants can be mistagged, usually at the beginning or ends of rows, when a stack of tags has terminated in the middle of a row and the next tag in the stack belongs elsewhere. Mistagging is usually spotted during preparation for photography or pollinations: the tags are easily moved, but their data must be checked. When tillers are confused with a clump of independent plants or plants have died before tagging, the mismatch between tags and plants prompts immediate reconsideration while standing in that row.

Another type of semantic problem arises from errors in data collection. Pollinations are the most vulnerable to scanning errors, since the same type of datum is scanned and assigning maternal and paternal identities depends on scanning the tags in the correct order. Our field procedure couples data collection tightly to pollination: often the pollinator stages stapling the tags to the bags with the data collector’s tag scanning, presenting each tag individually while naming the parents aloud. Similarly, the data collector reads back the phenotypes as the scoring data are collected, and the two resolve differences on the spot. Often two facts are collected: one for the original scoring and another for the contemporaneous scoring. Other semantic problems are detected only after data collection, usually by inspection. For example, selfing is planned for only certain rows and the plan is revised during the season as needed, so putative selves in other rows are checked against the current plan, the field, and eveyone’s memories. As *Chloe* is adapted to one’s wokflow, additional rules to detect one’s charateristic anomalies can be developed.

## 6 Discussion

The goal of *Chloe* is to be a flexible system that helps research groups of all sizes manage their work and the provenance of their data more efficiently. Unlike more targeted and integrated systems such as the Breeding Management System and Integrated Breeding Platform, *Chloe*’s code, equipment, and procedures make no assumptions about what experiment one is doing, the choice of data collection devices, or the uses to which the data will be put. In building *Chloe* and *Demeter*, I preferred platform flexibility, modularity, speedy execution, commodity hardware, and familiarity over custom graphical interfaces, rigid workflows, proprietary systems, and hiding data and ideas under too much machinery.

Since each research group values contemporaneous data collection and task automation differently, *Chloe*’s modular design and agnosticism let one choose which parts of one’s own workflows, equipment, and data warrant using or adapting *Chloe*. Many variations are possible.

- One could dispense entirely with barcoding, entering the data by hand into one’s current spreadsheets, exporting these to `csv` files, then pushing them into one’s own version of *Demeter* using local modifications of `convert_*_data.perl`.
- *Chloe* has several Perl scripts that produce tags not discussed here, such as plant tear-off, harvest, and row plan tags. For those who just want a barcode, say to identify a sample during photography, these scripts will be enough.
- We started with Symbol Technologies’ SPT 1800 barcode scanning devices, which displayed a data entry form the user designed on the device or on a PC using a proprietary application; checked each datum’s syntax and uniqueness as it was entered; and exported the collected data to a PC *via* a cable. The successors of these now obsolete scanners tend to be bulkier and more expensive, have shorter commercial lifetimes, and require specific computational platforms to run the proprietary configuration and synchronization apps. Instead, we opted for the commodity mobile devices, bluetooth scanners, spreadsheets, and cloud services when we had to replace our SPT 1800s.
- For those who prefer syntax checking at the moment of data entry, spreadsheet apps can now do some types of syntax checking and these functionalities are increasing.
- *Chloe* and *Demeter* accommodate pooling of pollen, ears, or seed, not just our normal individual pollinations of the first ear. Pollinating a divided ear with multiple males at the same time requires a minor extension only if it is important to indicate the orientation of the sectors.
- *Chloe*’s framework can easily be extended to other types of experiments and data, such as genomics and molecular biology; to other organisms, including prokaryotes; and to other types of equipment, including robots.

A group might focus on transforming their existing data into *Demeter*’s framework to simplify crop planning, pedigree construction, and field book generation. Others will find faster, semi-automated stock management more beneficial and dispense with plant tagging. These groups would just label their smallest filing group (for us, the virtual “sleeves”); enter the bounding seed packets of each group; revise the seed filing algorithm to match their inventory system; and then print packet labels for a quick checklist when packing seed for planting. Investigators just beginning their careers may want to adopt all of *Chloe* and *Demeter*, even if incrementally.

Placing downstream analyses in pipelines, rather than integrating them into the system, allows one to choose both the purposes of the analyses and the analytical approaches. For example, should it someday become important to use the camera parameters of the leaf images, I would write a rule that extracts them by calling ExifTool in the shell, parses the resulting file, and writes out a new type of fact. I would use exactly the same pipelining approach for other kinds of data and analyses, such as mapping, genotyping, genome-wide association, gene expression, proteomics, and biochemical studies, rather than reinvent specialist code.

Of necessity, *Chloe* includes some hard-wiring of syntax and semantics. This is a slight but real restriction on generalizability and requires some effort at adaptation. Examples of hard-wired syntax and semantics in *Chloe* include our conventions for numerical genotypes and other identifiers, the order of the columns in the data collection spreadsheets, and the rules that generate the field book. While in principle one could develop detectors for identifier syntax, one would need many working examples from many groups to develop something robust. Detecting the semantics of genetic experiments would be even more problematic, requiring long-term research in natural language processing to develop appropriate rules that capture an adequate domain model, or extensive data for machine learning, or both. Some types of data and tasks will be common among geneticists, so those parts should be widely useful and require little adaptation. Others will be specific to a laboratory or experiment: *Chloe* can be a model when designing these parts for your provenance and inventory systems. As much as possible, I have separated lab-specific syntax and organization into modules, or marked these as comments in the code, or both.

Two future tasks are to automate more quality checks in pedigree and genotype generation and to automate the generation of OWL 2-compliant metadata drawn from ontologies [10–12, 15]. The first task is straightforward, but the second is much more complicated. Like the public Genomes2Fields data [18], the semantics of *Chloe*’s data and computations are presented as structured comments in the files. What is left unsaid is any alignment of *Chloe*’s semantics with existing ontologies: to the best of my knowledge, current ontologies do not cover this sort of information at *Chloe*’s and *Demeter*’s level of granularity, though many existing ontologies express ideas that would be derived from work captured in *Chloe*. The ENVO environment ontology includes information on climates, habitats, soils, and environmental processes suitable for work in ecology and biodiversity, but these are very broad-brush descriptions compared to weather data or the USDA soil maps [3]. Ontologies for subsets of experimental procedures, such as the Ontology for Biomedical Investigations (OBI), Biological Imaging Methods Ontology (FBbi), Measurement Method Ontology (MMO), and Chemical Methods Ontology, cover specimen preparation for microscopy, clinical, or biochemical analyses [2, 13, 14, 16]. Similarly, ontologies for plant phenotypes (Phenotype and Trait Ontology (PATO), Plant Trait Ontology (TO), Plant Phenology Ontology (PPO), and Cotton Trait Ontology), genes and genomics (Planteome, reference 4), plant morphology (Plant Ontology (PO)), and agronomy (Agronomy Ontology (AGRO)) cover only portions of *Chloe* or are specialized for particular species [4, 5, 7]. Building a new ontology coordinated with these is a much more complicated task than simply generating metadata and must be community-driven to be genuinely useful. The impetus for such an effort could come when more groups develop experience with *Chloe* and want to share their raw, or near-raw, field data and methods.

## 7 Acknowledgments

I owe an enormous debt to my colleagues in the Missouri Maize Center. Ed Coe suggested I make all of my record-keeping electronic from the start, and this work is the direct result of that remark. Ed Coe, Gerry Neuffer, Georgia Davis, Theresa Musket, Kristen Leach (who modelled Figure 1, panel g), Karen Cone, Susan Melia-Hancock, and Sherry Flint-Garcia showed me the ropes of field work and their plant labelling and data collection methods. Jason Green and Jaturon Harnsomburana stuck out that first field season when we cobbled together the first pieces of what eventually became *Chloe*. Susan Melia-Hancock devised our tracking scheme for tassels, ears, and pollinations in the first year, which we use to this day, and Kristen Leach suggested the tablet/scanner pair. Gerry Neuffer showed me his provenance methods and especially, the principle of self-identification for every datum coming from the field, teaching by wonderful example. I had steady encouragement from Mary Schaeffer, Ed Coe, Jack Gardiner, Ramona Walls, and Carolyn Lawrence-Dill on ontologies and all things maize databases. Hope this work might be generally useful came from Carolyn Rasmussen, Damon Lisch, Tim Beissinger, and Jim Birchler. Rex Gish designed and built our plant tag fixture, and Ghasan Al Bahhash patiently saws our tags every year, despite my never bringing him the job as early as he would like. My own students over the years have stress-tested and bullet-proofed our procedures, contributing many improvements and helping me uncover silly assumptions I had baked into *Chloe*. Armani, Mac, and Vinny each contributed helpful discussions. This work was supported by NSF MCB-1122130, the MU Research Board, and an anonymous gift.

## Footnotes

1 For us, the only exception to this rule is sibbing wild types by mutants in mutant rows, where both parents have the same prefix. Here, we must look at the mutant phenotype data collected later to answer questions. We are extra vigilant during scanning when constructing double mutants, since that has no inherent preferred direction.

2 Syntax is the form of the data: the precise arrangement of characters a human or program uses to identify the type of a datum. Semantics is the meaning of the data: what a particular syntax denotes. For example, a date can be specified many different ways, such as 1/4/2020. Syntactically, we have *one digit slash one digit slash four digits*. Semantically, we have either *Gregorian day month four-digit-year* or *Gregorian month day four-digit-year*. Every computer language and application constrains and recognizes syntax. Recognizing and interpreting semantics remains a far harder task.

3 I am favoring geek vernacular here over more precise computational jargon: in Prolog, facts are a particular type of rule; the most general term is *predicate*.

4 A common example of extension is the annotation of genes by the Gene Ontology [1,6]: the annotation is individual to each gene because we don’t yet know the classification rules with sufficient accuracy.

